# Fgf signalling triggers an intrinsic mesodermal timer that determines the duration of limb patterning

**DOI:** 10.1101/2023.02.13.528267

**Authors:** Sofia Sedas Perez, Caitlin McQueen, Joseph Pickering, Kavitha Chinnaiya, Patricia Saiz-Lopez, Maria A. Ros, Matthew Towers

**Author notes:** Equal contribution (Sofia Sedas Perez, Caitlin McQueen).

## Abstract

Complex signalling between the apical ectodermal ridge (AER - a thickening of the distal epithelium) and the mesoderm controls limb patterning along the proximo-distal axis (humerus to digits). However, the essential requirement for AER-Fgf signalling during *in vivo* development makes it difficult to understand the exact roles that it fulfils. To overcome this barrier, we developed an amenable *ex vivo* chick wing tissue explant system that faithfully replicates *in vivo* parameters. Using inhibition experiments and RNA-sequencing, we identify a transient role for Fgfs in triggering the distal patterning phase. Fgfs are then dispensable for the maintenance of an intrinsic mesodermal transcriptome, which controls proliferation/differentiation timing and the duration of patterning. We also uncover additional roles for Fgf signalling in maintaining AER-related gene expression and in suppressing myogenesis. We describe a simple logic for limb patterning duration, which is potentially applicable to other systems, including the main body axis.

## Introduction

Much is known about how developing tissues and organs are spatially patterned. However, less is known about how the rate and/or duration of patterning events are determined both within and between different species (often referred to as developmental timing) ^1-4^. This is important because patterning events need to be temporally coordinated and larger species tend to develop at much slower rates than smaller species ^5^. The developing limb is an excellent system for understanding how the duration of patterning is determined as we have a deep knowledge of the underlying mechanism, which is composed of early and late patterning phases ^6^ (Figure 1a). The early proximal patterning phase (red) involves the stepwise activation of *Hoxa/d10/11* genes in proliferative distal mesoderm cells (dark blue circles in Figure 1a), and is considered to require the depletion of retinoic acid (RA) emanating from the main body of the embryo ^7-12^. Hoxa/d10/11 specify cells with proximal positional values, which instruct their development into the stylopod and the zeugopod ^13, 14^ (humerus and the ulna/radius - Figure 1a). The depletion of RA is influenced by growth of the limb away from the body and by the degradation enzyme, Cyp26b1, which is transcriptionally induced by opposing Fgf signals from the apical ectodermal ridge ^15, 16^ (AER-Fgfs – the AER is a thickening of the distal epithelium – Figure 1a). In different avian species (quail, chick and turkey), the duration of the early proximal patterning phase varies and takes between 12 and 30h ^17^. The loss of RA signalling in the distal part of the limb allows the initiation of the late distal patterning phase (light blue in Figure 1a) that coincides with *Hoxa/d13* gene activation ^11, 18^, which specifies the positional values of the autopod ^19^ (wrist and digits). In the quail, chick and turkey, the late patterning phase runs for a similar duration and takes between 48 and 54 h ^17^. Prolonging RA signalling in quail and chick wings extends the early patterning phase, and because the late phase remains unchanged, the duration of patterning is then equivalent to that of chick and turkey wings, respectively ^17^. It is likely that the degradation of RA and/or its transcriptional effectors (Meis1/2) is responsible for species’ differences in the duration of patterning.

**Figure 1.**
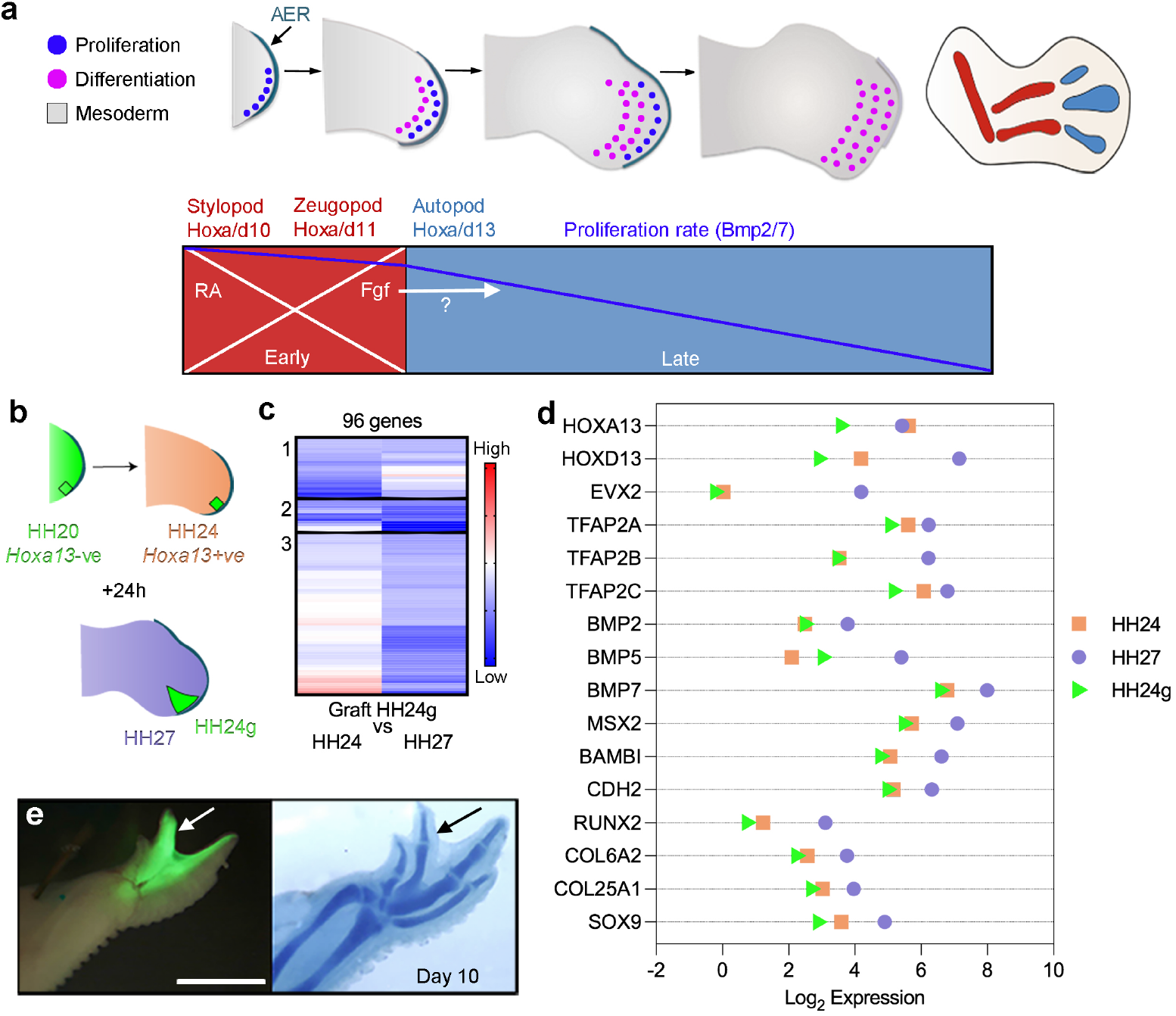
Proximo-distal patterning of the chick wing and the intrinsic mesoderm transcriptome. **a**) A population of undifferentiated proliferative distal mesoderm cells (dark blue circles) is maintained beneath the apical ectodermal ridge (AER) at the distal tip of the wing - cells displaced proximally differentiate (pink circles) into proximal structures and then distal structures as outgrowth proceeds (stylopod, zeugopod and then autopod). Early proximal patterning phase: retinoic acid (RA) signalling from the flank is opposed by Fgf signalling from the AER (AER-Fgfs), which permits the expression of *Hoxa/d10* and *Hoxa/d11* genes in the mesoderm that specify the positional values of the stylopod and the zeugopod. Late distal patterning phase: clearance of RA creates a permissive environment for the activation of *Hoxa/d13* genes in the mesoderm that specify the positional values of the autopod. An intrinsic mesodermal Bmp2/7-dependent timer is suggested to control the duration of the patterning phase, and the involvement of AER-Fgfs is an open question. **b**) Procedure to find the intrinsically-activated transcriptome: 150μm blocks of GFP-expressing (green) HH20 chick wing bud distal mesoderm was denuded of ectoderm and grafted under the AER of wild type HH24 buds (orange), incubated for 24h until the host stage is HH27 (purple) and the graft stage in HH24 (HH24g - green). **c**) Clustering of RNA-sequencing data across pairwise contrasts with the log2-fold change degree of gene expression indicated by the colour (red: higher, blue: lower). **d**) Plot showing expression levels of *Hoxa13, Hoxd13, Evx2, Tfapa, Tfapb, Tfapc, Bmp2, Bmp5, Bmp7, Msx2, Bambi, N-cadherin (Cdh2), Runx2, Col6a2, Col25a1* and *Sox9* as normalised log2 values of the RNA sequencing read-count intensities. **e**) Grafts of HH20 mesoderm made to HH24 buds often give rise to complete digits as shown at day 10 (*n*=12/35). Scale bar: 1mm.

During the late distal patterning phase the AER and underlying mesoderm continue to crosstalk via reciprocal signalling (epithelial-mesodermal or e-m signalling) ^20^. The AER is an essential structure and its removal in the chick wing truncates outgrowth ^21, 22^, which can be rescued by a bead soaked in Fgf protein implanted into the mesoderm ^23, 24^. In turn, the mesoderm produces the Bmp antagonist, Gremlin1 (Grem1), which maintains AER-Fgfs ^25, 26^. Embryological experiments performed on the chick wing suggest that an intrinsic Bmp2/7-dependent mesodermal proliferation timer controls the duration of the late distal patterning phase, which ends when all cells have differentiated ^18, 27, 28^ (pink circles in Figure 1a). The role of AER-Fgfs in this model is to maintain mesoderm proliferation as permissive factors (Figure 1a). By contrast, mouse genetics support an instructive role for AER-Fgfs during the late distal patterning phase ^29-34^. However, the essential requirement for AER-Fgfs in limb development makes it difficult to understand the roles that they fulfil ^31^.

Here, we use heterochronic mesodermal grafting techniques coupled with RNA-sequencing to further decipher the extent to which the late distal patterning phase is controlled by extrinsic or intrinsic mechanisms. We then develop an experimentally amenable chick wing tissue explant system to examine the roles of Fgfs. We provide evidence that Fgfs are required for initiating the late distal patterning phase by activating *Hox13* genes. However, they are not required thereafter for intrinsic mesodermal proliferation/differentiation timing and in determining the duration of patterning. We also reveal unexpected roles for Fgfs in maintaining AER-related gene expression and in suppressing myogenesis.

## Results

### The intrinsic chick wing mesoderm transcriptome

To understand the extent to which the late distal patterning phase is intrinsically regulated in the mesoderm we identified the underlying transcriptome. Previous heterochronic grafting techniques revealed that *Hoxa13* is intrinsically-activated in chick wing mesoderm according to the age of the donor tissue, and that proliferation parameters, as well as cell adhesion properties are also maintained ^18^. To identify the intrinsic mesodermal transcriptome, we used the same approach coupled with RNA-sequencing. We grafted 150μm blocks of HH20 (Hamburger Hamilton) chick wing distal mesoderm beneath the AER of HH24 host wing buds, and allowed them to develop for 24h, so that the developmental stage of the graft is HH24g (graft) and the host HH27 (Figure 1b).

After HH20 grafts were made to HH24 wings and left for 24h, we performed RNA– sequencing on 150μm of the distal-most grafted mesoderm, and for controls, equivalent (non-grafted) tissue in the contralateral wing bud. This identified 303 differentially expressed genes: 73 between HH24 and HH24g datasets and 230 between HH27 and HH24g datasets (>2-fold difference with an adjusted *p*-value of < 0.05 – Supplementary data 1 and Supplementary data 2). We then used hierarchical clustering analyses to identify those genes that are intrinsically-activated in the grafts like *Hoxa13*. Based on our previous analyses, we expected intrinsically-activated genes to be expressed in the grafts (HH24g) at lower levels when compared with host levels (HH27), consistent with their later activation ^18^. Three out of nine clusters contain genes that behave in this manner: cluster 1 (23 genes), cluster 2 (12 genes) and cluster 3 (61 genes (Figure 1c; Supplementary figures 1-3). Expression was also reduced in the grafts (HH24g) compared to donor levels (HH24), suggesting that the grafting procedure slightly delays gene activation as we noticed previously ^18^. However, the majority of the genes show similar trends in plots of the read-counts of the RNA-seq data, indicating that expression in HH24g samples (green triangles) is generally lower than, or equivalent to HH24 samples (orange squares), but substantially lower than HH27 samples (purple circles - Figure 1d). The identification of *Hoxa13* confirmed the results of our previous study ^18^ and is the only gene characterised in limb development that is found in cluster 1. *Hoxd13* and *Evx2*, which are co-regulated syntenic genes, are found in cluster 2. Many genes that are involved in limb development are present in cluster 3, including three that encode members of the Ap2 family of transcription factors, *Tfap2a, Tfap2b* and *Tfap2c*, which have been implicated in maintaining the undifferentiated state of the distal mesoderm ^35, 36^. In addition, several genes that encode members of the Bmp pathway are found, including the *Bmp2, Bmp5* and *Bmp7* ligands, and their downstream transcriptional effector, *Msx2. Bambi* is also present, which encodes a pseudoreceptor that limits the range of Bmp signalling in chick wing distal mesoderm ^37^. These findings confirm our earlier study implicating Bmp signalling as a central determinant of an intrinsic proliferation timer in the mesoderm ^28^. Additional genes of interest in cluster 3 include *N-cadherin*, which encodes a molecule that mediates cell adhesion along the proximo-distal axis ^38^, and differentiation regulators, including *Col25a1* and *Col6a2* (connective tissue), *Sox9* (cartilage) and *Runx2* (bone). This is consistent with the finding that grafts of HH20 distal mesoderm made to HH24 buds often give rise to a complete digit (Figure 1e). We did not identify genes that could act downstream of Bmp signalling in progressively suppressing G1 to S-phase entry, possibly because core cell cycle regulators are predominantly controlled at the post-translational level. However, a likely candidate is the Cyclin D inhibitor, *p57*^*kip2* 39^, which we found is transcriptionally induced 24h after Bmp2-soaked beads implanted into the distal mesoderm of HH24 wings (Supplementary figure 4). These data reveal the transcriptome involved in intrinsic mesoderm development.

### Chick wing explants proliferate with *in vivo* rates

Genes associated with Fgf signalling were not identified in the intrinsic mesoderm transcriptome (Supplementary figures 1-3). However, it is possible that Fgf signalling permissively maintains intrinsic gene expression. This is difficult to validate experimentally because of the essential requirement for the AER and *Fgfs* for *in vivo* limb development ^21, 22, 31^. We investigated whether a tissue explant system could circumvent the essential functions of the AER and Fgfs, by culturing the posterior-distal third of HH20 (day 3.5 to 4 of incubation) chick wing buds in Matrigel (Figure 2a). We included the Sonic hedgehog (Shh) producing polarising region that is found in the posterior-distal mesoderm, and which makes a reciprocal signalling loop with AER-Fgfs ^40, 41^. We determined whether *in vivo* proliferation parameters are maintained in explants using flow cytomtery, which gives an accurate stage-specific read-out of cell cycle rate ^18, 39^ (percentage of cells in G1-phase). The analyses reveal that there is no significant difference in proliferation rates between *in vivo* tissue and explanted tissue at 24 and 48h (Figure 2b). In addition, EdU labelling shows that cells are actively progressing into S-phase at 48h (Figure 2c). Furthermore, lysotracker staining does not reveal any appreciable levels of apoptosis at 48h (Figure 2d). We also determined gene expression parameters using the multiplex hybridisation chain reaction (HCR) *in situ* technique followed by light-sheet microscopy. Over time, *Meis1* and *Hoxa11* expression are excluded from the distal mesoderm and become restricted to proximal regions in both wing buds and explants (Figures 3a-d, note, distal is the top of the panels and posterior is the right-hand side). In addition, *Hoxa13, Hoxd13, Shh, Bmp2, Sox9* and *Runx2* are expressed in distal regions of both wing buds and explants (Figures 3e-p). In explants, there is a slight delay of approximately 6-12h in the clearance of *Meis1* and *Hoxa11* from distal regions, and in the activation of *Hoxa13* and *Runx2*, which is likely to be caused by acclimatisation to the culturing conditions (Figures 3a-d, e-f, o-p). Digit condensations are observed in wing buds as indicated by *Sox9* and *Runx2* expression at 72h (Figures 3m and o) but they are not as pronounced in explants (arrowheads - Figures 3n and p). Taken together, these findings indicate that chick wing bud explants closely replicate *in vivo* parameters.

**Figure 2.**
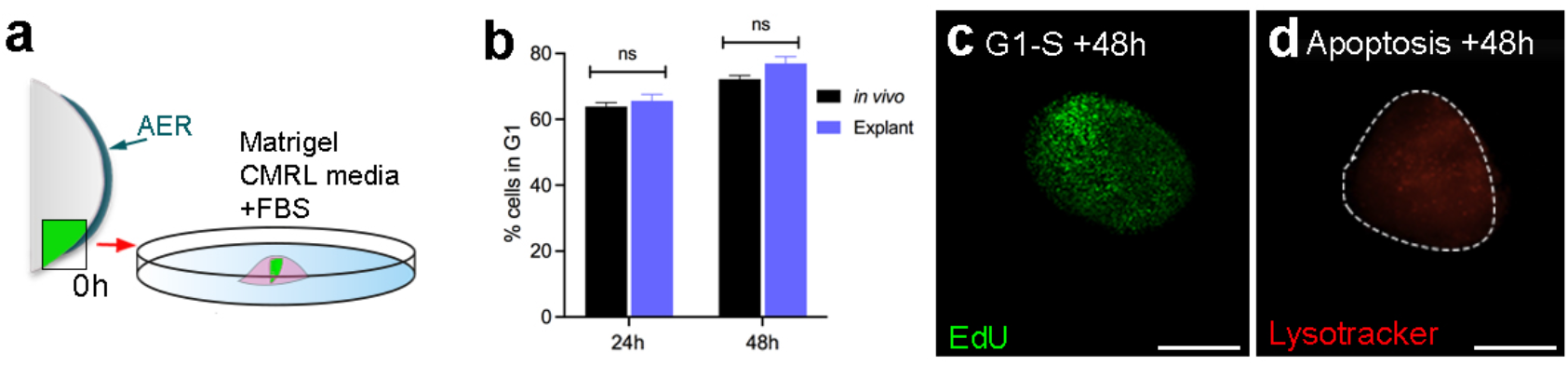
Proliferation timing in chick wing explants. **a**) The posterior-distal region of the wing bud including the AER and the polarising region is dissected at HH20 (designated as 0h) and cultured in Matrigel, CMRL media and FBS. **b**) Flow cytometric analyses at 24 and 48h reveals no significant difference in the percentage of cells in G1-phase in explants and equivalent *in vivo* tissue: two-tailed unpaired *t*-tests were carried out at each time-point (*p-*valu*e* >0.05 – bars indicate standard error). **c**) EdU labelling shows that cells are progressing into S-phase in explants (*n*=4/4). **d**) Lysotracker staining shows no appreciable apoptosis in explants (*n*=16/16). Scale bars in c and d – 200μM.

**Figure 3.**
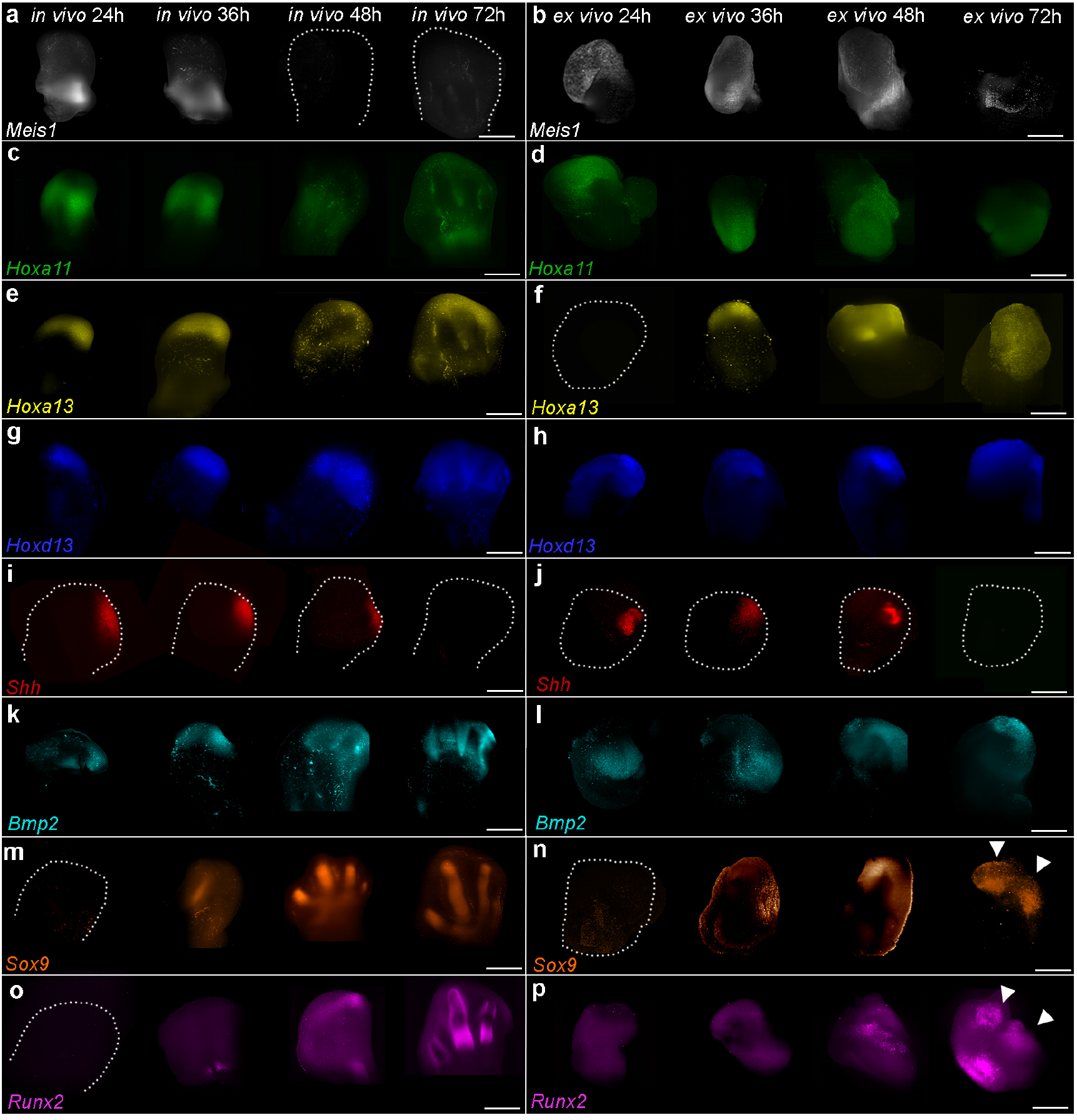
Gene expression timing in explants. **a, b**) *Meis1*, **c, d**) *Hoxa11*, **e, f**) *Hoxa13*, **g, h**) *Hoxd13*, **i, j**) *Shh*, **k, l**) *Bmp2*, **m, n**) *Sox9* and **o, p**) *Runx2* expression in wing buds (*in vivo*) and explants (*ex vivo*) over 72h shown by HCR in situ hybridisation (*n*=>3 in all cases - proximal is the bottom of the panels; posterior is the right). There is a delay in the clearance of *Meis1* (**a, b**) from the distal part of explants and in the activation of *Hoxa13* (**e, f**). Digit condensations are marked by *Sox9* and *Runx2* expression in (**m, o**) wing buds and (**n, p**) explants (arrowheads). Scale bars for wing buds - 200μM and explants - 500μM.

### Chick wing explants denuded of AER proliferate with *in vivo* rates

To begin to test the requirement of Fgfs in the late distal patterning phase, we removed the AER at 0h, as shown by the loss of *Fgf8* expression at 24h (Figures 4a and b). In addition, *Shh* expression is also undetectable, consistent with a role for the AER in maintaining the activity of the polarising region ^40-42^ (Figures 4a and b). The removal of the AER dramatically attenuates Fgf signalling, as determined by a downstream readout, *Mkp3* (also known as *Dusp6* and *Pyst1*) ^43^, which is either absent or expressed at very low levels at 24h (Figures 4c and d). This finding indicates that signalling by mesodermal Fgfs (including Fgf10 that reciprocally maintains Fgf8 activity ^44^) is also affected by the removal of the AER. Unexpectedly, despite losing the activity of both the AER and the polarising region, flow cytometric analyses reveal no significant difference in the percentage of cells in G1-phase of the cell cycle in explants compared with controls at 24 and 48h (Figure 4e). Additionally, EdU labelling demonstrates that cells are transiting from G1 to S-phase at 24h (Figures 4f and g). Furthermore, although explants cultured without an AER appear to have a smaller surface area, it is not significantly different to explants cultured with an intact AER (Figure 4h). Lysotracker staining indicates that apoptosis is enhanced in the posterior necrotic zone in explants cultured without the AER at 24h, but it is not enhanced in other regions (Figures 4i, j). These findings reveal that the AER and the polarising region are dispensable for mesodermal proliferation in explants.

**Figure 4.**
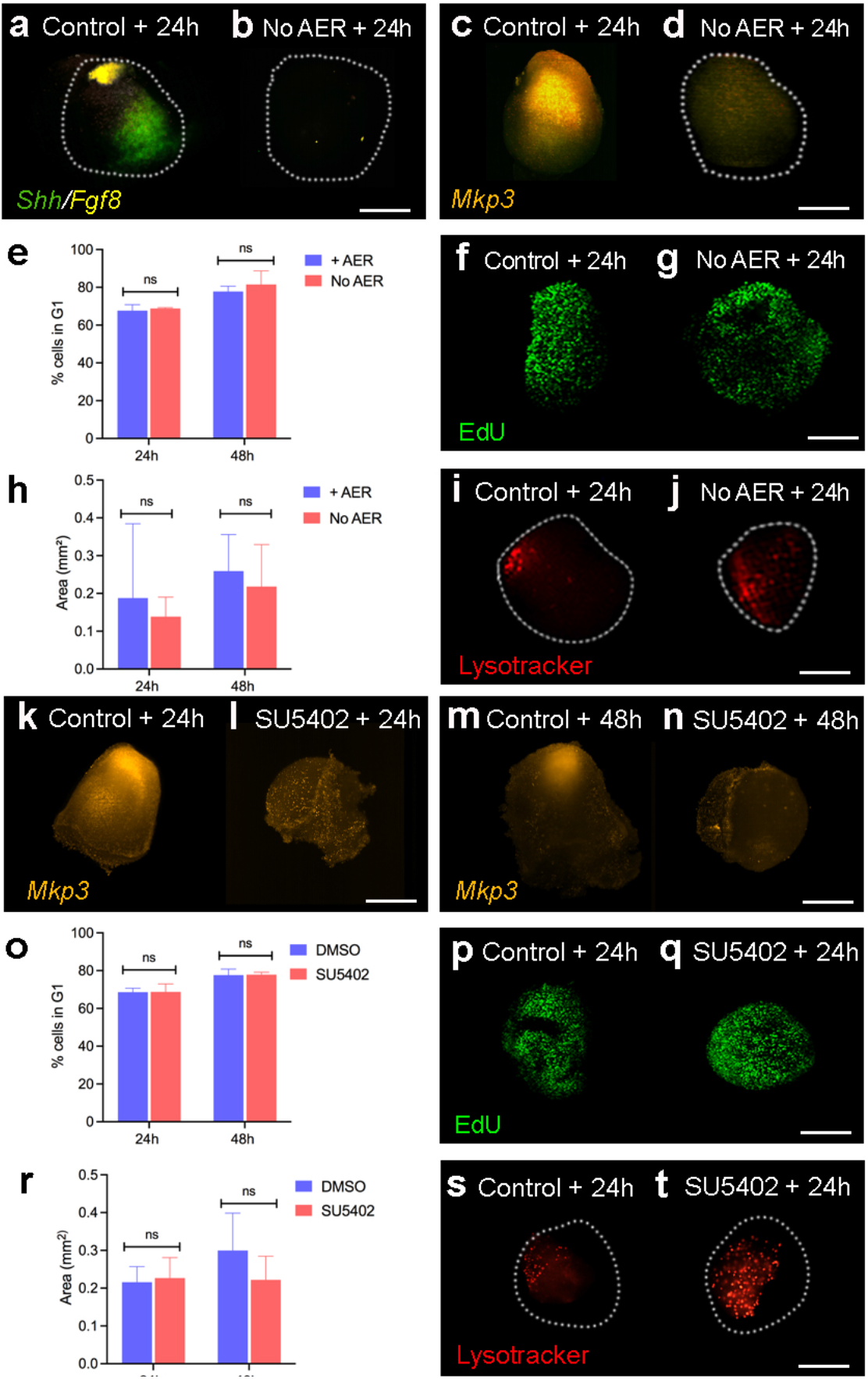
AER removal and Fgf signalling inhibition in explants. **a, b**) *Shh* and *Fgf8* are undetectable 24h after AER removal (*n*=4/4). **c, d**) *Mkp3* is severely down-regulated 24h after AER removal (*n*=3/3). **e**) Flow cytometric analyses reveal no significant difference in the percentage of cells in G1-phase in explants cultured with or without the AER at 24 and 48h: two-tailed unpaired *t*-tests were carried out at each time point (*p-*valu*e* >0.05 – bars indicate standard error). **f, g**) EdU labelling reveals cells are progressing into S-phase in explants cultured with or without the AER at 24h (*n*=3/3). **h**) The surface area of explants is not significantly different between explants cultured with or without the AER at 24 and 48h: two-tailed unpaired *t*-tests were carried out at each time point (*n*=>30, *p-*value >0.05 – bars indicate standard deviation). **i, j**) Lysotracker staining reveals apoptosis in the posterior necrotic zone, which is slightly increased in explants cultured without the AER at 24h (*n*=6/6). **k**-**n**) *Mkp3* expression is severely down-regulated 24 and 48h after 5μm of SU5402 is added to the media (*n*=8/8 in both experiments). **o**) Flow cytometric analyses reveal no significant difference in the percentage of cells in G1-phase between explants cultured with control DMSO or SU5402 at 24 and 48h: two-tailed unpaired *t*-tests were carried out at each time point: (*p-*value >0.05 - bars indicate standard error). **p, q**) EdU labelling demonstrates that cells are progressing into S-phase in explants cultured with DMSO or SU5402 at 24h (*n*=9/9). **r**) The surface area of explants is not significantly different between explants cultured with DMSO or SU5402 at 24 and 48h: two-tailed unpaired *t*-tests were carried out at each time point (*n*=7; *p-*value >0.05 - bars indicate standard deviation). **s, t**) Lysotracker staining detects apoptosis in the posterior necrotic zone, which is increased in explants cultured SU5402 at 24h (*n*=12/12). Scale bars - 200μM.

### Chick wing explants proliferate with *in vivo* rates when Fgf signalling is attenuated

To inhibit Fgf signalling in chick wing explants in a more systematic way than by removing the AER, we used a pharmacological approach by applying the Fgf receptor 1 inhibitor, SU5402, to the culture medium at 0h. *Mkp3* is observed at low/background levels in explants treated with SU5402 at 24 and 48h (Figures 4k-n). In addition, no significant difference in the percentage of cells in G1-phase of the cell cycle is observed in explants treated with or without SU5402 for 24 and 48h, as determined by flow cytometry (Figure 4o). Furthermore, EdU labelling confirms that cells are actively progressing from G1 to S-phase at 24h (Figures 4p, q). Explants treated with SU5402 have a similar surface area compared to controls at 24h, and appear smaller at 48h, although the difference is not significant (Figure 4r). Consistent with this observation, lysotracker staining indicates increased apoptosis in explants treated with SU5402 for 24h when compared with controls (Figures 4s, t). Therefore, the attenuation of Fgf signalling and the removal of the AER have similar effects on chick wing explant development, and neither is required for mesodermal proliferation timing.

### Fgf signalling is required for 5’ *Hox13* expression and for maintaining the AER and the polarising region

We sought to determine the global requirement of Fgf signalling during the distal patterning phase, by adding SU5402 to the culture medium, and then performing RNA-sequencing at 48h on entire explants. In total, the expression of 1600 protein coding genes is significantly affected by SU5402 application, of which, 937 are up-regulated and 663 are down-regulated (2x-fold change with an adjusted *p*-value of <0.05 - Supplementary data 3). We concentrated on genes that have known roles in Fgf signalling and/or limb patterning. All four *Sprouty* (*Spry*) genes, which are targets of Fgf signalling ^45^ are down-regulated, with no detectable expression of *Spry3* (Figure 5a - note that log_2_-fold changes are shown as well as actual read-counts). In addition, *Mkp3* is strongly reduced (Figure 5a), consistent with the HCR *in situ* data (Figures 4m, n). *Fgf8* is undetectable, and several other genes expressed in the AER are strongly down-regulated, including *Sp8, Hoxc13* and *Dlx5/6* ^46^ (Figure 5a). Consistent with the AER removal experiments, *Shh* is completely abolished, and downstream effectors of the signalling pathway, including *Hhip, Ptch1/2* and *Gli1* ^47^ are significantly down-regulated (Figure 5a). *Hoxa13* is undetectable, *Hoxd13* and *Evx2* are significantly down-regulated, and *Hoxa11*/*d10/11/12* are moderately reduced (Figure 5a). Expression of *Bmp2* and its downstream targets *Msx1* and *Sox9* are also slightly reduced ^48^ (Figure 5a). Very few genes known to be involved in limb patterning are up-regulated, but exceptions include *Grem1*, which encodes the AER maintenance factor, and *Alx4*, which represses *Shh* expression ^49^. HCR *in situ* hybridisation for *Hoxa13, Hoxd13, Shh, Fgf8, Bmp2* and *Sox9* at 48h (Figures 5b-m) confirms the RNA-sequencing data (Figure 5a). These observations reveal critical roles for Fgf signalling in *Hoxa/d13* activation, and in maintaining the AER and the polarising region. The general stability of *Bmp2* and *Sox9* expression suggests that Fgf signalling is dispensable for the onset of chondrogenesis.

**Figure 5.**
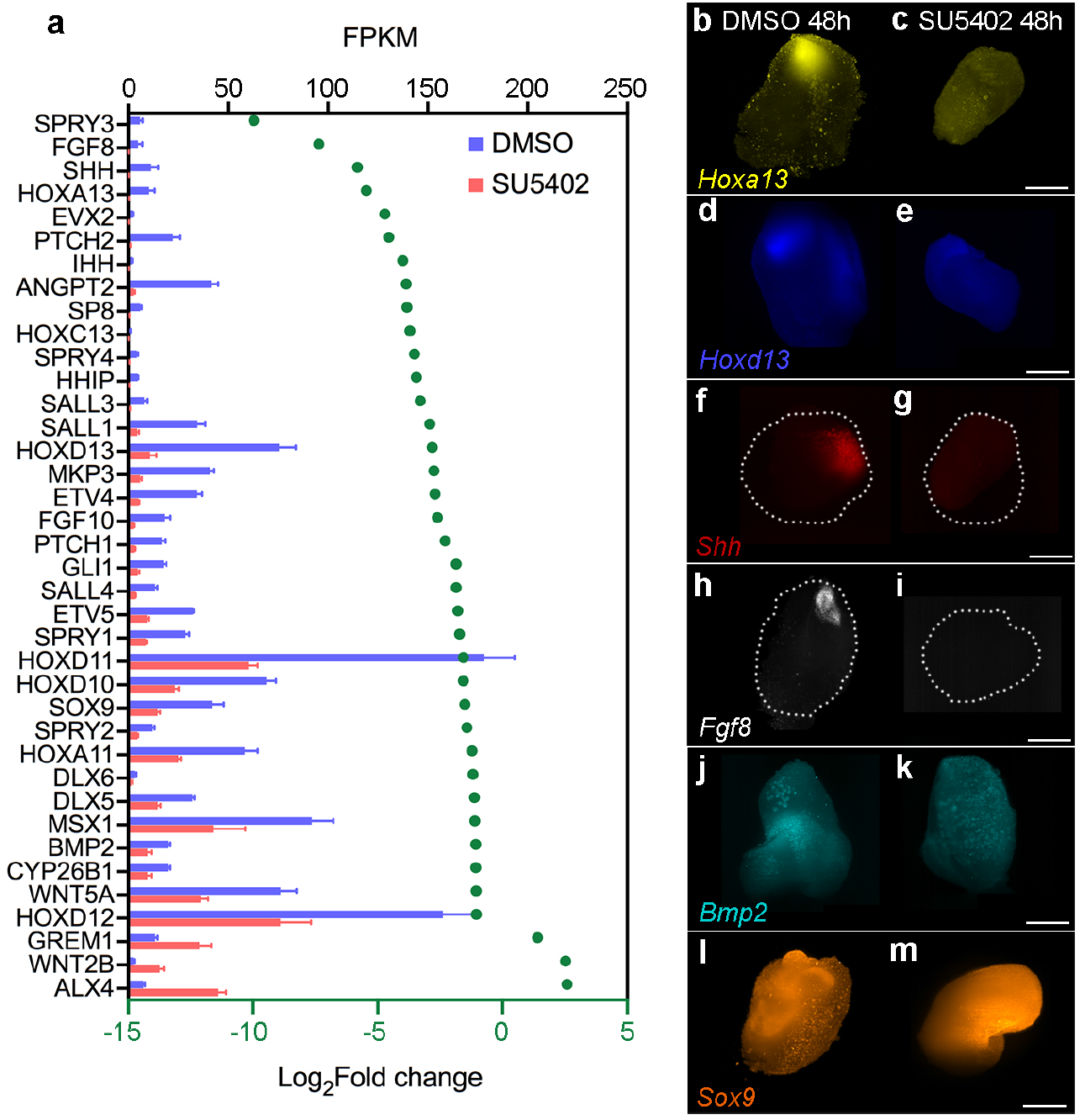
RNA sequencing of SU5402-treated explants. **a**) Expression of genes involved in Fgf signalling and/or limb development in control DMSO- and SU5402-treated explants at 48h as shown by log2-fold changes (>2-fold change; adjusted *p*-value of <0.05) and normalised read-counts mapped to each gene (FKPM – Fragments of Kilobase of Exon Per Million Mapped Fragment - bars indicate standard deviation). **b, c**) *Hoxa13*, **d, e**) *Hoxd13*, **f, g**) *Shh*, **h, I** *Fgf8*, **j, k**) *Bmp2* and **I, m**) *Sox9* expression are down-regulated in SU5402-compared with DMSO-treated explants as shown by HCR *in situ* hybridisation at 48h (*n*=4/4 in each example). Scale bars - 200μM

### Requirement for Fgf signalling in maintaining the intrinsic mesoderm transcriptome

We then analysed how the stability of the intrinsic mesoderm transcriptome (Figure 1) is affected by the attenuation of Fgf signalling in explants (Figure 5). Analysis of the RNA-sequencing data reveals that only 15% of intrinsically-activated genes are significantly down-regulated and 5% are significantly up-regulated (Figure 6 - 2x-fold change with an adjusted *p*-value <0.05). Notably, *Hoxa13*, and *Hoxd13* and *Evx2* are among the five most down-regulated genes, the others being *Acot12* and *Sez6l*, which have not been characterised in limb development. Other notable genes that are slightly down-regulated include *Bmp2, Bmp7* and *Sox9*. These data show that Fgf signalling is required for the expression of *Hox13* genes, but not for many of the other intrinsically-activated genes.

**Figure 6.**
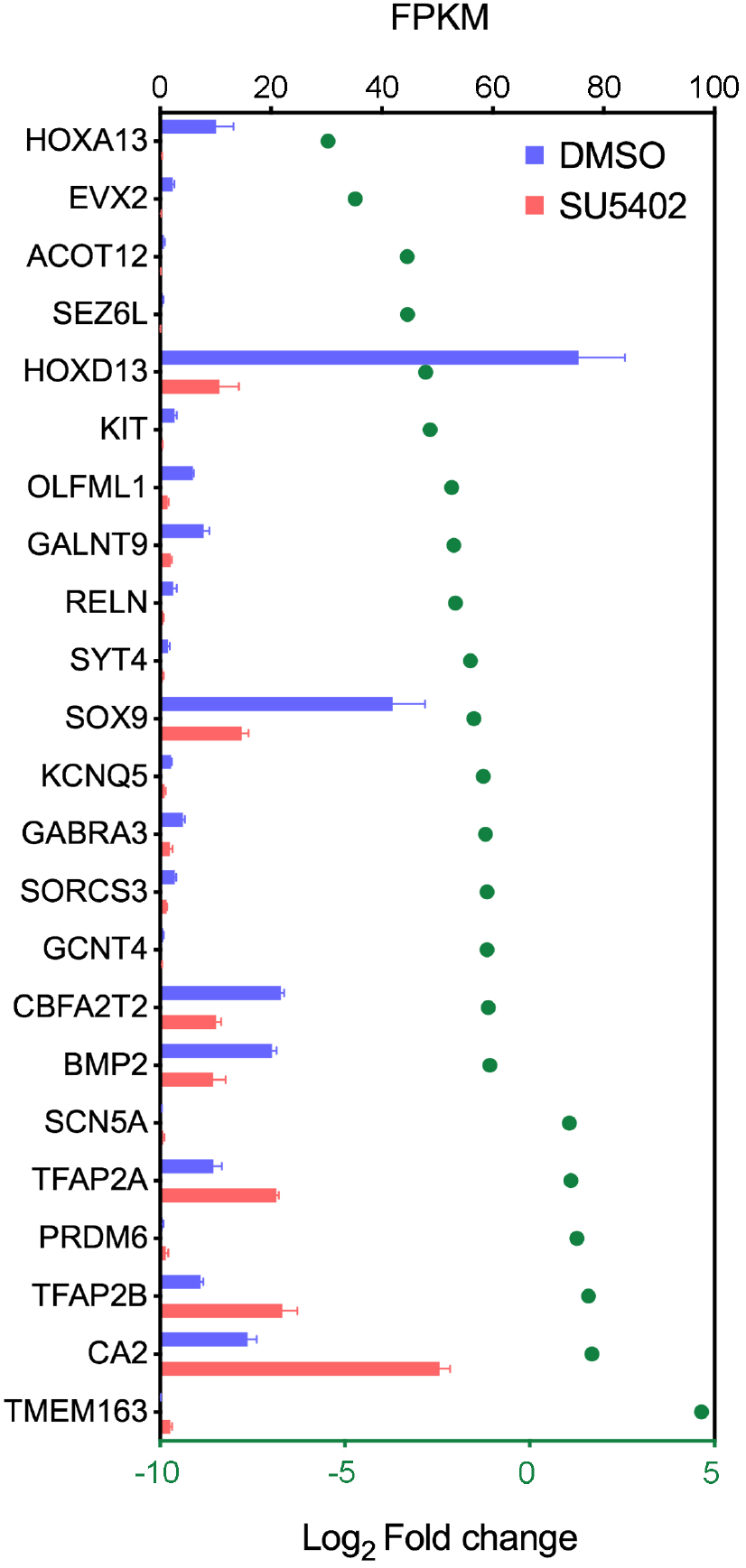
Requirement of Fgf signalling for intrinsic mesodermal gene expression. Intrinsically-expressed genes (Figure 1) that show a significant change in expression when Fgf signalling is attenuated in explants (Figure 5) as shown by log2-fold changes (>2-fold change; adjusted *p*-value of <0.05) and read-counts mapped to each gene (FKPM – Fragments of Kilobase of Exon Per Million Mapped Fragment – bars indicate standard deviation).

### Fgf signalling suppresses myogenesis

Unexpectedly, the RNA-sequencing data reveal that the inhibition of Fgf signalling in explants causes a significant up-regulation of genes representing all steps of the myogenic pathway at 48h (Figure 7a – Supplementary data 3). These genes include *Pax3* and *Pax7*, which are markers of uncommitted myogenic precursor cells; *Myf5* and *MyoD1*, which are involved in myogenic commitment; *Myogenin* (*MyoG*), which regulates myogenic induction, and several *Myosin light chain* (*Myl*) and *Myosin heavy chain* (*Myh*) genes, which are involved in myogenic differentiation ^50^ (Figure 7a). *Pax3*-positive myogenic progenitor cells migrate into the limb from the somites ^50^, and HCR *in situ* hybridisation reveals a substantial population in tissue dissected at 0h to make explants, thus explaining why myogenic gene expression is detected (Figure 7b – *Shh* and *Fgf8* are also shown). *Pax3, Myf5, Myod1* and *Myog* are restricted to dorsal muscle masses at 48h in normal wing development (Figures 7c, f, i, l) and are detectable in control explants (Figure 7d, g, j, m). However, SU5402 treatment causes their significant up-regulation in explants (Figures 7e, h, k, n), consistent with the RNA-sequencing data (Figure 7a). These findings show that attenuation of Fgf signalling enhances myogenic gene expression in explants.

**Figure 7.**
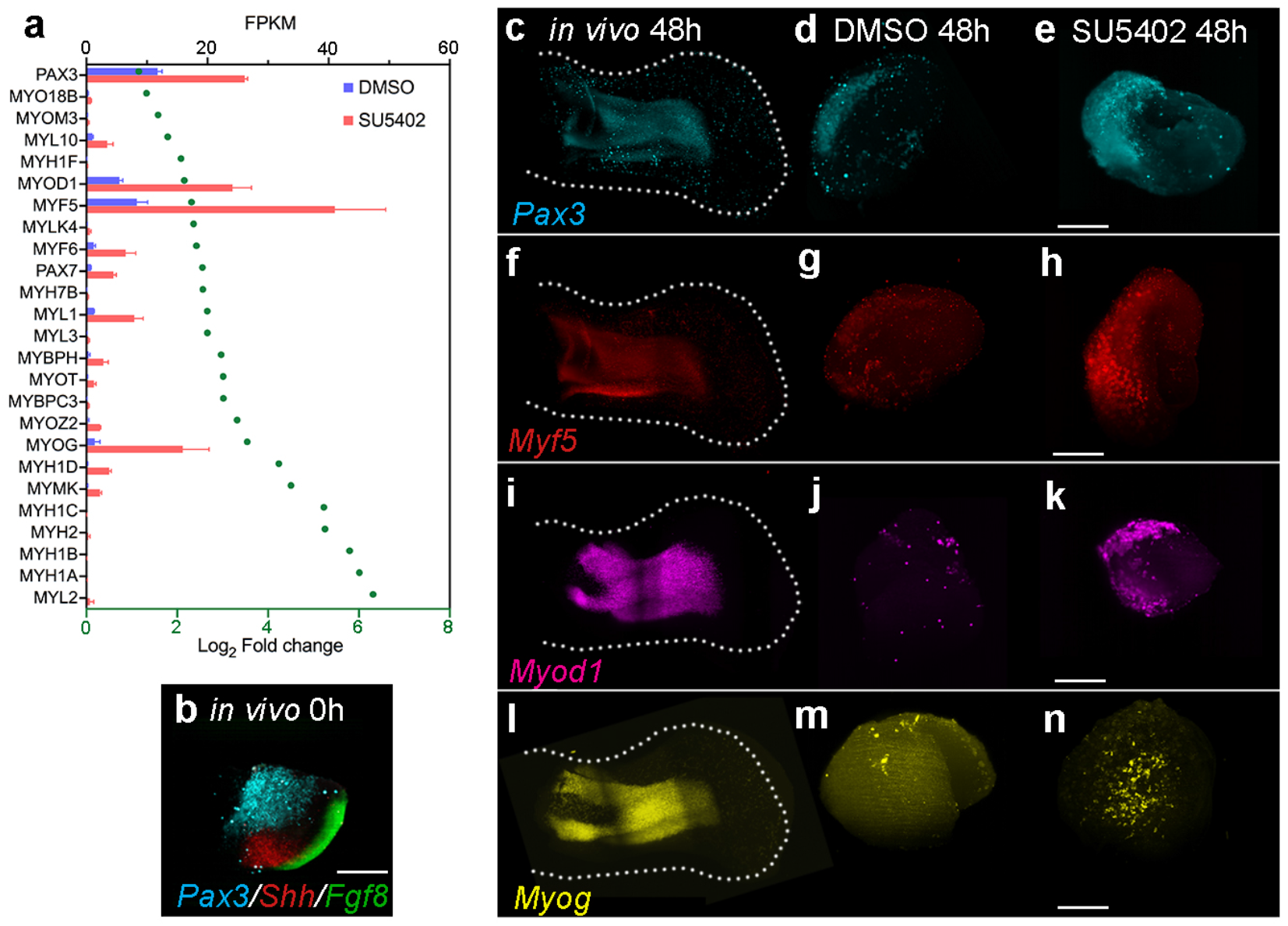
Myogenic gene expression in SU5402-treated explants. **a**) Expression of genes involved in myogenesis in control DMSO- and SU5402-treated explants at 48h as shown by log2-fold changes (>2-fold change; adjusted *p*-value of <0.05) and normalised read-counts mapped to each gene (FKPM – Fragments of Kilobase of Exon Per Million Mapped Fragment - bars indicate standard deviation). **b**) *Pax3*-positive myogenic progenitor cells are present in regions of the wing bud used to make explants as shown by HCR *in situ* hybridisation at 0h (*Shh* and *Fgf8* also shown *n*=9/9). **c**) *Pax3*, **f**) *Myf5*, **i**) *Myod1* and **l**) *Myog* expression in wing buds at 48h (*n*=>4). **d, e**) *Pax3*, **g, h**) *Myf5*, **j, k**) *Myod1* and **m, n** *Myog*, are up-regulated in SU5402 compared with DMSO-treated explants at 48h (*n*=7/7 in each example). Scale bars - 500μM in wings; 200μM in explants.

## Discussion

We have developed a chick wing tissue explant system to investigate the role that Fgf signalling fulfils in limb patterning. We demonstrated that Fgf signalling is required for initiating the late distal patterning phase of limb development by activating *Hoxa/d13* genes. Following this critical role, we unexpectedly revealed that Fgf signalling is dispensable for the mesoderm to maintain its intrinsic patterning parameters. In addition to maintaining the AER through the Fgf10-Fgf8 feedback loop, we uncovered an additional novel role for Fgf signalling in suppressing myogenesis.

### Limb patterning duration

The antagonism between proximal signals (considered to be RA) and distal signals (AER-Fgfs) controls the early (extrinsic) patterning phase characterised by the expression of *Meis1/2* genes (*Meis*), and the specification of the stylopod and zeugopod ^7-12^ (Figure 8). Genetic analyses in the mouse limb indicate that AER-Fgfs induce the expression of the RA-degrading enzyme, *Cyp26b1* ^15^, whose product creates a gradient of RA signalling, and therefore of Meis ^12^. In this model, Meis levels need to be low enough to allow the transition from stylopod to zeugopod specification (Figure 8). Indeed, the level of RA signalling correlates with—and can change the rate of—5’ *Hox* activation between different avian species ^17^.

**Figure 8.**
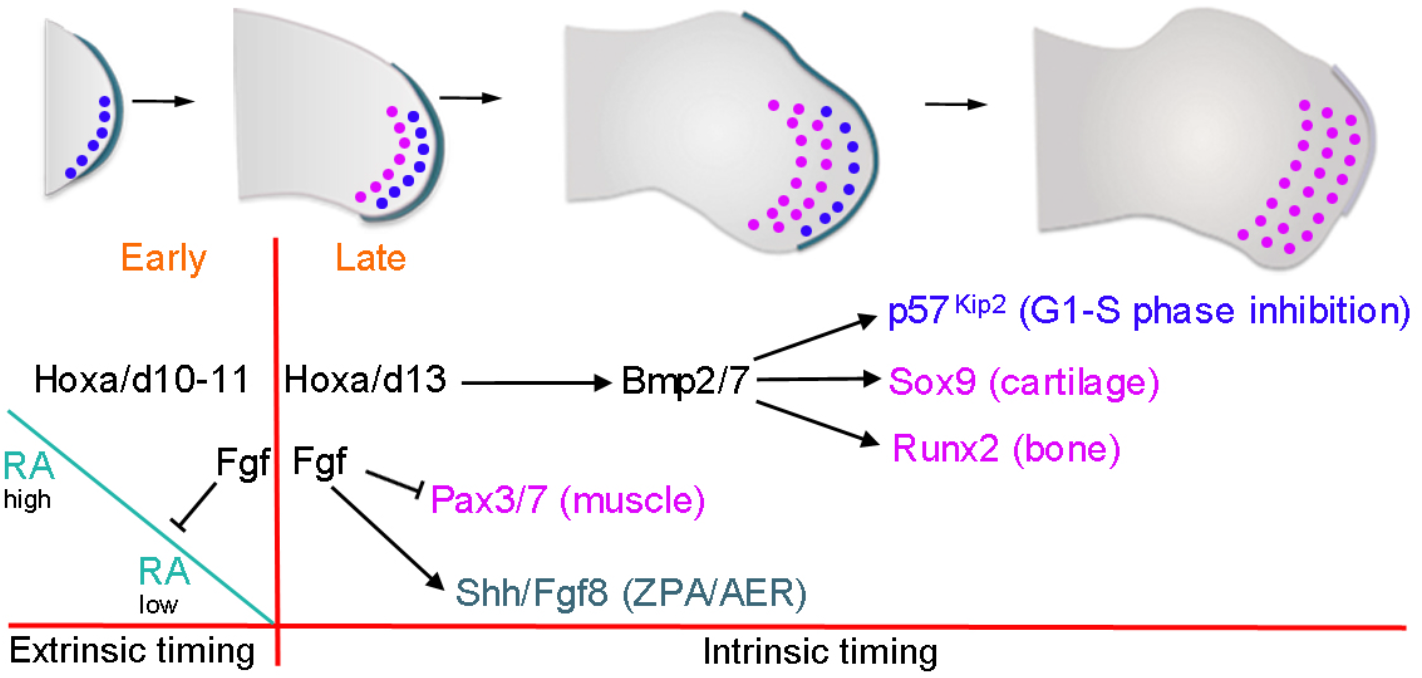
Model of chick wing patterning duration. Early proximal extrinsic patterning phase - antagonistic flank-derived Retinoic acid (RA) and AER-derived Fgf extrinsic signals time *Hoxa/d10*/*11* gene activation that specify the positional values of the stylopod and the zeugopod. Late distal intrinsic patterning phase - Fgf-depletion creates a permissive environment for the activation of *Hoxa*/*d13* genes that specify the positional values of the autopod. Hoxa/d13 activates *Bmp2/7* independently of Fgf signalling to intrinsically control the timing, hence the duration of mesodermal proliferation, via activation of *p57*^*kip2*^, which inhibits G1-S-phase progression, and *Sox9/Runx2*, which promote chondrogenesis/osteogenesis. Fgf signalling suppresses myogenesis and maintains the polarising region (ZPA-*Shh*) and the AER (*Fgf8*), which are permissively required for outgrowth.

Evidence from both the mouse and chick suggest that the instructive AER-Fgf dependent clearance of RA (Meis) from the distal part of the limb creates a permissive environment that is required for *Hoxa13* expression ^8-12^ and for the activation of the late (intrinsic) distal patterning phase (autopod specification) ^18^. Removal of the AER in the chick wing causes the immediate loss of *Hoxa13* expression, which can be restored with an Fgf-soaked bead ^51^. Once *Hoxa/d13* genes are activated, the distal mesoderm gains intrinsic properties including a Bmp-dependent proliferation timer that progressively inhibits G1 to S-phase entry and determines the duration of chick wing patterning (Figure 8) ^28^. By developing an *ex vivo* chick wing tissue explant system, we unexpectedly revealed that distal mesoderm cells maintain normal proliferation/differentiation timing when Fgf signalling is severely curtailed. Under these conditions, our data indicate that *Hoxd13* is activated at sufficient levels to initiate the late distal patterning phase in the absence of Hoxa13. These observations support genetic analyses in the mouse, demonstrating that distal development is relatively normal in *hoxa13*^-/+^/*hoxd13*^-/-^ compound forelimbs (note distal development fails in *hoxa13*^-/-^/*hoxd13*^-/-^ limbs) ^19^. Similarly, Fgf8 can support distal development in the mouse limb in the absence of the other AER-Fgfs - Fgf4, Fgf9 and Fgf17 ^31^. Our results indicate that, although *Hox10*/*11* genes are down-regulated, they are relatively stable after Fgf signalling is inhibited, consistent with results showing that they are still expressed following AER removal in the chick wing ^52^. Therefore, Fgf signalling is unexpectedly dispensable for the intrinsic distal programme once it is activated.

The intrinsically-activated transcriptome allows us to propose a basic gene regulatory network (GRN) that underlies the late distal patterning phase (Figure 8). Previous work showed that Hoxa13 directly activates the expression of *Bmp2* and *Bmp7* in distal regions of the mouse limb ^53^, thus providing a mechanism that coordinates proliferation and differentiation (Figure 8). We presented evidence that Bmp signalling regulates the Cyclin D-dependant kinase inhibitor, *p57*^*kip2*^, which could control the decline in the rate of proliferation in the distal mesoderm (Figure 8). In addition, Bmp signalling induces differentiation by activating primary regulators of chondrogenesis (*Sox9*) and osteogenesis (*Runx2*) ^54^ (Figure 8). It is striking that the application of either Bmp2 or Bmp7 partially rescues the phenotype of *hoxa13*^-/-^ mutant mouse limbs ^53^, thus demonstrating the pivotal nature of this pathway during the late patterning phase. Therefore, an implication of our findings is that the Fgf-dependent activation of *Hoxa/d13* determines the duration of patterning, by triggering an intrinsic self-terminating process based on the Bmp-dependent timing of proliferation and differentiation (Figure 8). Other intrinsically-activated genes that are likely to act downstream of Hoxa/d13 include the cell adhesion molecule, *N-cadherin*, which could provide distal mesoderm cells with different positional values (e.g., phalange vs. carpal) ^38^. It is notable that Bmp signalling progressively increases in the limb ^28^, and also induces its own inhibitors, such as *Grem1* ^55^. Therefore, it is likely that the dynamics of key effectors of the Bmp signalling pathway determine the duration of the late distal patterning phase.

Our results prompt a re-evaluation of the direct functions of Fgf signalling in limb development. We found that the attenuation of Fgf signalling causes the loss of *Shh* expression in the polarising region, consistent with previous reports ^40, 41^, but also loss of AER-expressed genes including *Fgf8*, which is probably due to the interruption of the Fgf10-Fgf8 feedback loop ^32, 33, 44^. Therefore, a minimal proliferation/differentiation timing GRN can operate without the two principle organisers of limb development (Figure 8). This could appear surprising because previous work described important roles for Shh signalling in controlling proliferation ^39, 56^. However, our results here indicate that Fgf signalling is required for supporting the mitogenic functions of Shh signalling. Indeed, the ability of Shh signalling to modulate polarising region proliferation dynamics requires an overlying AER ^39^. It is likely that the function of the AER during *in vivo* limb development includes a direct mechanical role that is dispensable in explants. Thus, Fgfs are required for the dorso-ventral flattening of the limb and outgrowth away from the main body ^57^, possibly by controlling planar cell polarity ^58, 59, 60^ and directional cell division ^61^. Loss of such processes could contribute to the excessive apoptosis that occurs when the AER is removed ^29^ or when Fgf signalling is genetically ablated ^31^.

Our results also suggest that Fgfs are not required for the maintenance of an undifferentiated progenitor state in the distal mesoderm, which according to the maintained expression of the *Tfap* family of transcription factors in our heterochronic mesoderm grafts, is an intrinsic property. We also revealed that Fgf signalling suppresses the myogenic pathway: a role that we uncovered because normal proliferation/differentiation trajectories are maintained in explants when Fgf signalling is attenuated (Figure 8). These data support early findings in which the over-expression of *Fgf4* in the chick wing inhibited myogenesis ^62^. Therefore, the explant system uncovers a minimal GRN that orchestrates the differentiation of the three primary classes of limb tissue - cartilage, bone and muscle (Figure 8).

### Perspectives

Parallels can be drawn with the patterning of the limb and of the main body axis. Neuromesodermal progenitors (Nmps) located at the posterior end of the main body transition from producing anterior axial structures - including the vertebrae of the trunk - to producing the posterior tail bud. This involves a switch in gene regulatory activity in which Gdf11 suppresses the anterior programme by inducing the RA-degrading enzyme, *Cyp26a1* ^63^, to activating the posterior programme via *Hoxb/c13* ^64^ - a process reminiscent of the function of Fgf signalling in switching proximal to distal limb patterning by antagonising RA signalling via *Cyp26b1*, which allows *Hoxa/d13* activation ^15^. During limb development, this switch involves *Hoxd* gene transcription directed by regulatory regions in topologically associating domains (TADs) located 3’ (up to *Hoxd11*) and 5’ (*Hoxd13*) to the cluster ^65^. It is unclear if distinct TADs govern the transition from trunk to tail patterning in the main body axis ^66^, and if this involves a switch from an extrinsic to an intrinsically-regulated programme that we describe here for limb development. We speculate that a common intrinsic programme exists downstream of *Hox13* activation that determines the duration of limb and tail patterning. This could be revealed with heterochronic grafting/RNA-sequencing experiments on chick Nmps, similar to those that we have performed in the limb.

## Materials and methods

### Chick husbandry and tissue grafting

Wild type and GFP-expressing Bovan Brown chicken eggs (Henry Stewart, Norfolk, UK) were incubated and staged according to Hamburger Hamilton ^67^. For tissue grafting, a 150μm strip of distal mesoderm, including the AER, was dissected from HH20 wing buds. The epithelium was removed after incubation in 0.25% trypsin at room temperature for 2 mins and the mesoderm was then cut into cubes, which were placed in slits between the AER and underlying mesoderm in the mid-distal region of HH24 wing buds using a fine sharpened tungsten needle.

### Chick wing tissue explants

The posterior third of HH20 wing buds were dissected in ice-cold PBS under a LeicaMZ16F microscope using a fine surgical knife. A bed of Growth Factor Reduced Matrigel (Corning) was prepared and allowed to set in four-well plates for 30-40 min at 37°C. The explants were then placed on top of the Matrigel (4-5 per well), covered with another layer of Matrigel, and cultured in CMRL media supplemented with 10% FBS/1% Pen Strep/1% L-Glut in a humidified incubator with 5% CO2 at 37°C. Explants were collected by replacing the culture media with Cell Recovery Solution (Corning) on ice.

### AER removal

The AER of wing buds of HH20 embryos was stained with 1% Nile blue solution *in ovo* and removed by pulling with sharp forceps. Embryos were then collected and the right-hand wing bud dissected to make explants as described above (the left-hand wing bud was used to make control explants).

### SU5402 treatment

SU5402 (Sigma) dissolved in DMSO with a final concentration of 5uM was added to the explant culture media at 0h.

### Explant size measurements

Explants were placed in a Petri dish containing 1XPBS and imaged using a LeicaMZ16F microscope. The surface areas of the explants were measured using the Record Measurement Feature of the Lasso tool in Adobe Photoshop. The measurement scale was set based on the metadata of the images.

### Hybridisation chain reaction *in situ*s

Samples were fixed in 4% PFA at 4°C overnight, then washed in PBS and progressively dehydrated through a methanol series and stored in methanol at -20°C. The samples were then rehydrated in a methanol to PBT series and treated with proteinase K for 5-7 min for explants and 15-20 min for wing buds, followed by post-fixing in 4% PFA for 20 min. At this point an optional bleaching protocol described below was performed. Samples were further washed with PBT, 5X SSCT and Molecular Instruments Probe hybridisation buffer before addition of the probe overnight at 37°C. The probes were prepared by adding 8 μl of 1μM probe to 500 μl of Molecular Instruments Probe hybridisation buffer. The next day samples were washed with Molecular Instruments Probe Wash Buffer, 5X SSCT and Molecular Instruments Amplification buffer. Amplifier pairs were prepared by heat-shocking at 95°C for 90 sec and snap cooling for 30 min in the dark at room temperature, before being added to the samples in the Molecular Instruments Amplification buffer. The samples with amplifiers in Molecular Instruments Amplification buffer were incubated overnight in the dark at room temperature. The next day samples were washed and stored in 5X SSCT before imaging. Fluorescent images of HCR and EdU labelling were taken with a Zeiss Z1 Lightsheet Microscope with a 10X objective. Images were processed with ImageJ (FIJI) and Adobe Photoshop. Limb bud images in Figure 7 were taken as tiled images and stitched together with the Grid/Collection stitching plugin in ImageJ (FIJI) ^68^. Autofluorescence bleaching was used for HCR on the limb buds in Figure 7. Samples were washed with PBT followed by incubation in a 3% H2O2 20mM NaOH PBT solution on ice for 3h to reduce sample autofluorescence. The samples were then washed in PBT for 3×10 min 3 before proceeding with HCR.

### Whole mount *in situ* hybridisation

Embryos were fixed in 4% PFA overnight at 4°C, dehydrated in methanol overnight at -20°C, rehydrated through a methanol/PBS series, washed in PBS, then treated with proteinase K for 20 mins (10μg/ml^-1^), washed in PBS, fixed for 30 mins in 4% PFA at room temperature and then prehybridised at 69°C for 2 h (50% formamide/50% 2x SSC). 1μg of antisense DIG-labelled mRNA probes were added in 1ml of hybridisation buffer (50% formamide/50% 2x SSC) at 69°C overnight. Embryos were washed twice in hybridisation buffer, twice in 50:50 hybridisation buffer and MAB buffer, and then twice in MAB buffer, before being transferred to blocking buffer (2% blocking reagent 20% lamb serum in MAB buffer) for 2 h at room temperature. Embryos were transferred to blocking buffer containing anti-digoxigenin antibody (1:2000) at 4°C overnight, then washed in MAB buffer overnight before being transferred to NTM buffer containing NBT/BCIP and mRNA distribution visualised using a LeicaMZ16F microscope.

### Bead implantation

Affigel beads were soaked in human Bmp2 protein (0.05μg/μl^1^ - R&D) dissolved in PBS/4mM HCl. Beads were soaked for 2 h and implanted into distal mesoderm using a sharp needle.

### Flow cytometry for cell cycle analyses

Explants and equivalent regions of stage-matched distal mesoderm were dissected in ice cold PBS under a LeicaMZ16F microscope using a fine surgical knife and pooled from replicate experiments (*n*=10-12), before being digested into single cell suspensions with trypsin (0.5%, Gibco) for 30 mins at room temperature. Cells were briefly washed twice in PBS, fixed in 70% ethanol overnight, washed in PBS and re-suspended in PBS containing 0.1% Triton X-100, 50μg/ml^-1^ of propidium iodide and 50μg/ml^-1^ of RNase A (Sigma). Dissociated cells were left at room temperature for 20 mins, cell aggregates were removed by filtration and single cells analysed for DNA content with a FACSCalibur flow cytometer and FlowJo software (Tree star Inc). Based on ploidy values cells were assigned in G1, S, or G2/M phases and this was expressed as a percentage of the total cell number (5,000-12,000 cells in each case). Statistical significance of numbers of cells in different phases of the cell cycle (G1 vs. S, G2 and M) between pools of dissected wing bud tissue and explants (12-15 in each pool) was determined by two-tailed unpaired *t*-tests to obtain *p*-values (significantly different being a *p*-value of less than 0.05).

### EdU labelling

Explants were incubated in 0.5mM EdU in CMRL for 4 hours at 37°C after release from Matrigel, then fixed with 4% PFA for 15 min and washed with 3% BSA/PBS at room temperature, before being permeabilized with 0.5% Triton X-100 in PBS for 20 min and washed with 3% BSA/PBS. Explants were incubated in the dark for 1 h in a click-kit reaction cocktail containing Azide Dye (Molecular Probes). This was followed by 3 washes in 3% BSA/PBS before imaging.

### Apoptosis assays

Explants were transferred to a Lysotracker (Life Technologies, L-7528)/PBS solution (1:1000) in the dark, incubated for 1 h at 37°C, washed in PBS, and fixed overnight in 4% PFA at 4°C. Explants were then washed in PBS and progressively dehydrated in a methanol series before imaging.

### RNA sequencing analyses and clustering

RNA sequencing was performed on two conditions: grafts as described above with equivalent *in vivo* tissue from the contralateral wing, and on explants treated with either DMSO or SU5402. Three replicate experiments were performed for each condition (*n*= 10-12 tissue samples in each experiment). Samples were collected by flash freezing in dry ice. Total RNA was extracted from samples using Trizol-chloroform extractions. Explant RNA was further concentrated using Zymo RNA clean and concentrator kit as per manufacturer’s instructions. Messenger RNA was purified from total RNA using poly-T oligo attached magnetic beads for explants. Sequencing was performed in either Illumina HiSeq 2000 PE50 (Grafts) or Illumina NovaSeq 6000 PE150 (Explants). Sequencing data were mapped using HISAT v2.0.3 (Grafts) or v2.0.5 (Explants) to the chicken reference genome. Raw data have been deposited in array express (https://www.ebi.ac.uk/arrayexpress/experiments/E-MTAB-6437/) for grafts and on GEO (GSE223444) for explants. Based on quality control checks one of the HH24 and HH24g samples was excluded from further analysis. Quantification of gene expression level for Explants was performed with Feature Counts v1.5.0-p3 and then FPKM (Fragments Per Kilobase of transcript sequence per Millions base pairs sequenced) of each gene was calculated based on the length of the gene and read counts mapped to this gene. Differential expression analysis between DMSO and SU5402 samples was performed using DESEq2 v1.20.0. Genes with an adjusted *p*-value <0.05 and a fold-change >2 were assigned as differentially-expressed. For grafts, the count data for the samples were normalised using trimmed mean of *m*-value normalisation and transformed with Voom, resulting in log2-counts per million with associated precision weights. A heat-map was made showing the correlation (Pearson) of the normalised data collapsed to the mean expression per group. A statistical analysis using an adjusted *p*-value < 0.05 and a fold-change >2 identified differentially expressed genes in the two contrasts evaluated. Gene clusters were identified from the set of differentially expressed genes. The evaluation considered between two and 35 clusters using hierarchical, *k*-means, and PAM clustering methods based on the internal, stability and biological metrics provided from the clValid R package. Most of the internal validation and stability metrics indicated that either the lowest possible number or conversely the highest number evaluated were preferable. The metrics giving more nuanced information in the intermediate range were the Silhouette measure, and the Biological Homogeneity Index (BHI). Based on manual inspection it was decided to use hierarchical clustering with *k*=9 gene clusters, which showed favourable properties for both these measures.

### Alcian blue skeletal preparations

Embryos were fixed in 90% ethanol for 2 days then transferred to 0.1% alcian blue in 80% ethanol/20% acetic acid for 1 day, before being cleared in 1% KOH.

## Acknowledgements

This work was supported by the Wellcome Trust (202756/Z/16/Z) P) to MT and by the Spanish Ministry of Science and Innovation (Grant PID2020-114525GB-I00) to M.R. We thank Marysia Placzek for critical reading.

