## Supplementary material for "Fgf signalling triggers an intrinsic mesodermal timer that determines the duration of limb patterning": Supplemetary figure

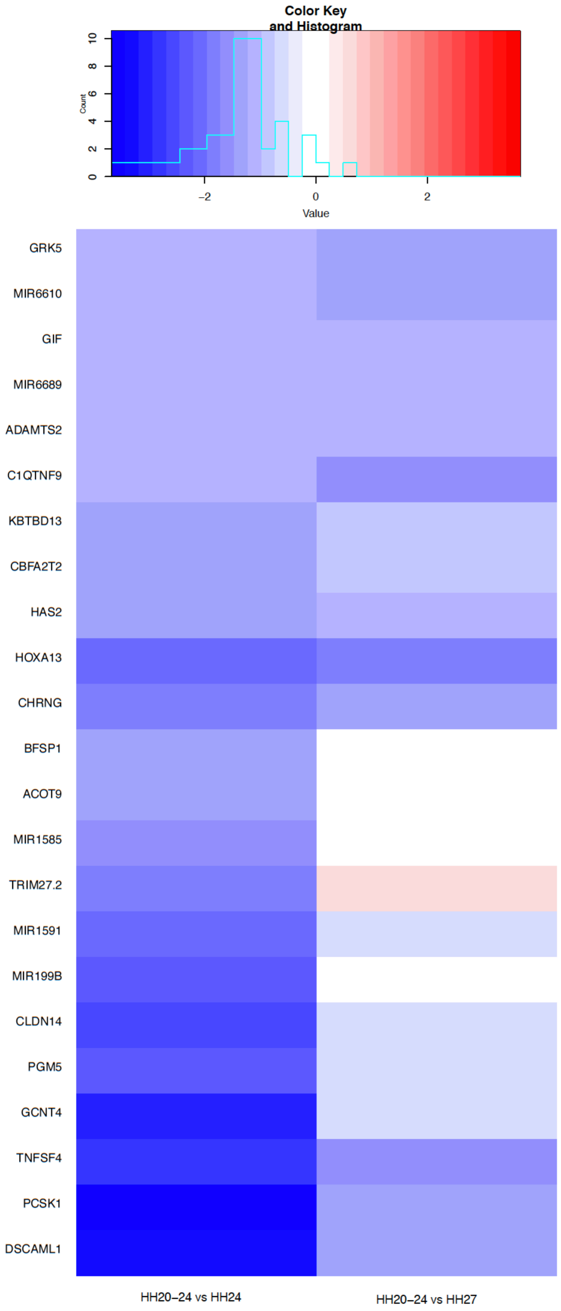


**Figure 1: Cluster 1**
23 genes including *Hoxa13* generally show much lower expression in stage HH20-24 grafts (HH24g) after 24h compared with donor stage HH24 tissue and also host stage tissue HH27.

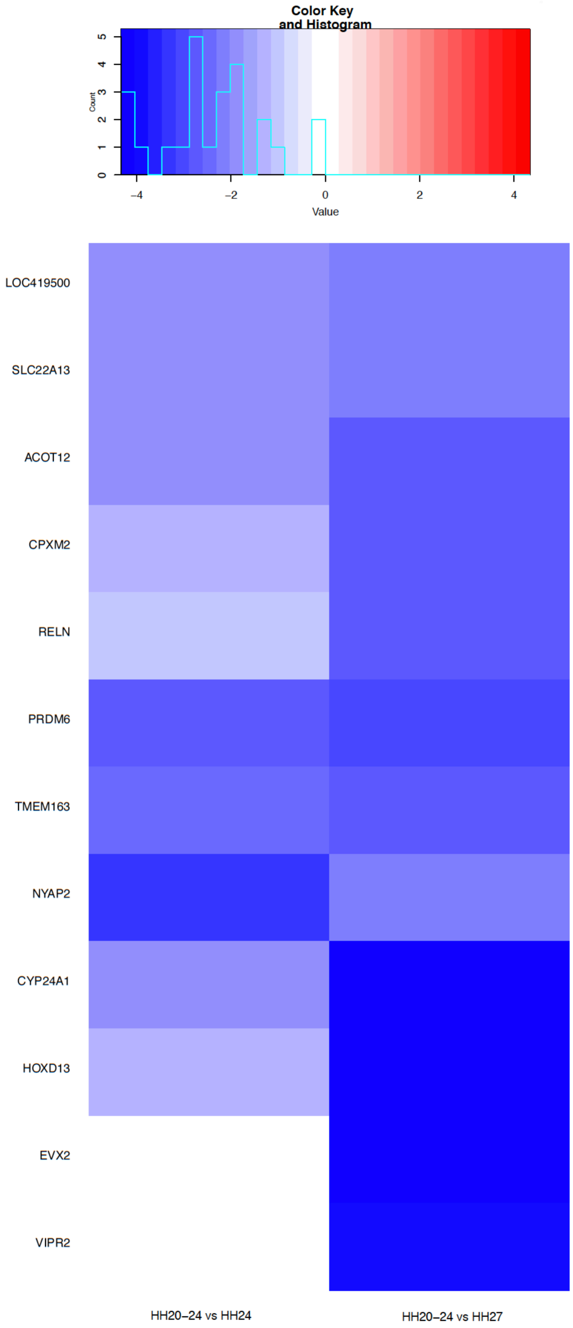


**Figure 2: Cluster 2**
12 genes including *Hoxd13* generally show much lower expression in stage HH20-24 grafts (HH24g) after 24h compared with donor stage HH24 tissue and also host stage tissue HH27.

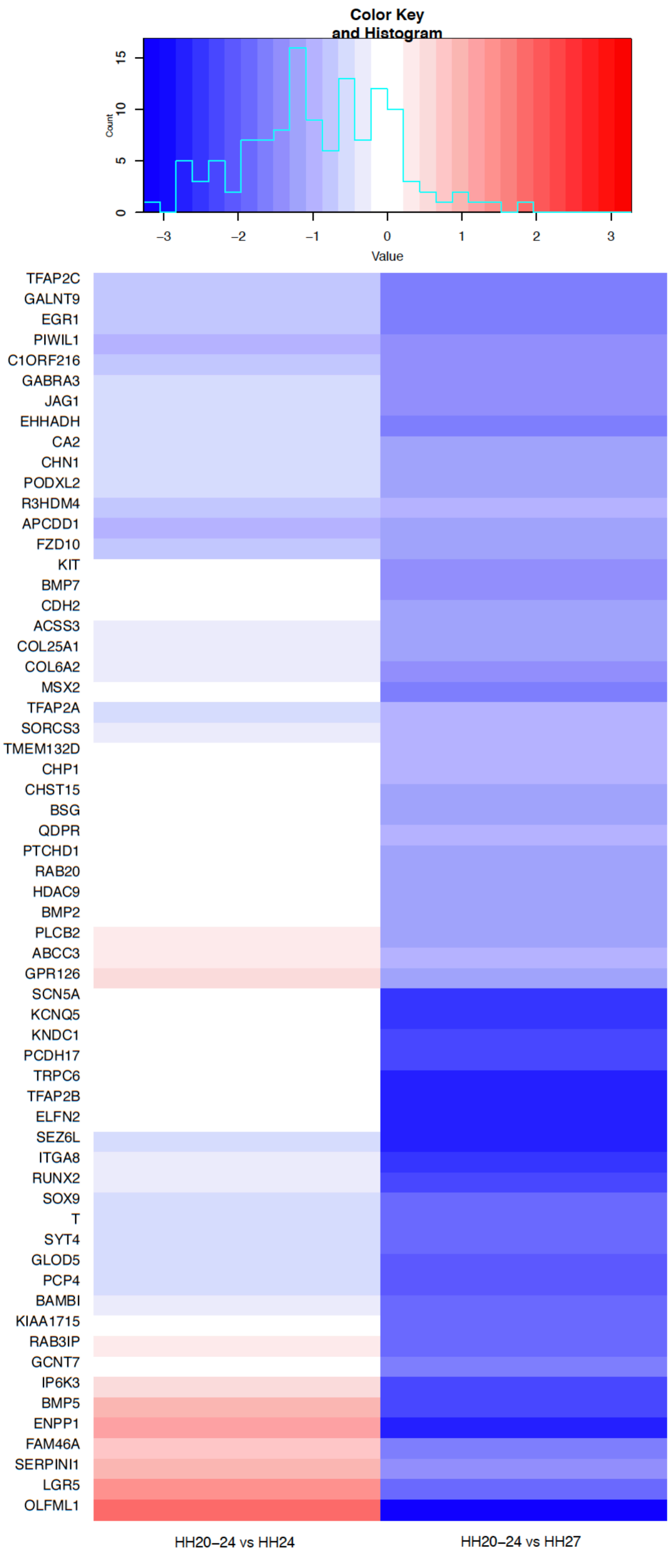


**Figure 3: Cluster 3**
61 genes including several associated with Bmp signalling and differentiation generally show similar levels of expression in stage HH20-24 grafts (HH24g) after 24h compared with donor stage HH24 tissue, and much lower expression compared with host stage tissue HH27.

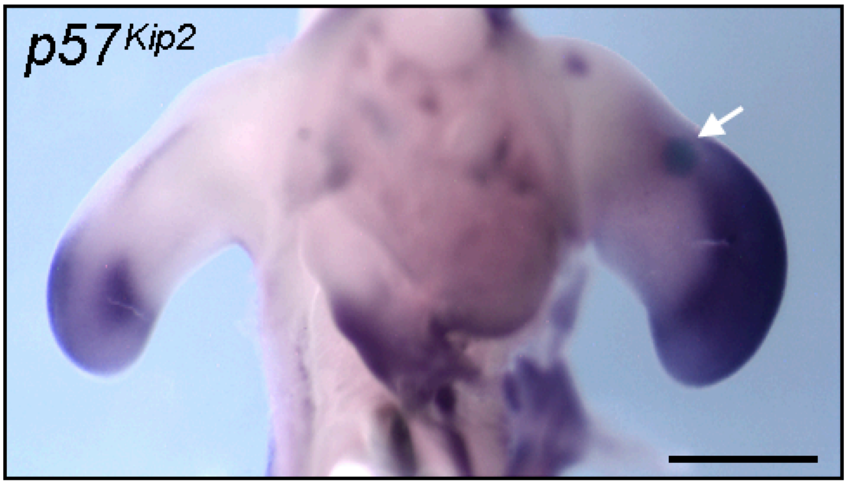


**Figure 4: Bmp2 regulation of *p57$^{kip2}$***

Bmp2 soaked beads (white arrow) implanted into the distal mesoderm of HH24 wing buds up-regulate *p57$^{kip2}$* expression 24h later at HH27 (*n*=4/4). Scale bar - 500μm
